# Mapping human cerebral blood flow with high-density, multi-channel speckle contrast optical spectroscopy

**DOI:** 10.1101/2025.03.03.638332

**Authors:** Byungchan (Kenny) Kim, Alexander C. Howard, Tom Y. Cheng, Jessica E. Anderson, Bernhard B. Zimmermann, Eric Hazen, Laura Carlton, Mitchell Robinson, Marco Renna, Meryem A. Yucel, Stefan A. Carp, Maria Angela Franceschini, David A. Boas, Xiaojun Cheng

## Abstract

Recently, speckle contrast optical spectroscopy (SCOS) enabled non-invasive, high signal-to-noise-ratio (SNR) human cerebral blood flow (CBF) measurements, relevant for both neuroscience and clinical monitoring of diseases with CBF dysregulation. Single channel SCOS measurements limit the information obtained to only one location on the head. In this work, we develop a multi-channel SCOS system to map spatial heterogeneity in CBF changes during human brain activation. Using a galvanometer, we temporally multiplexed a free-space laser to 7 source fibers positioned at different locations on the head. Diffuse light collected from the tissue is captured by fiber bundles projecting to 17 complementary metal-oxide semiconductor (CMOS) cameras, resulting in 50 source-detector channels measuring optical density (OD) and relative CBF changes covering an area of 7.6 cm by 6.6 cm on the head. We validated the spatial specificity and stability of the system using a liquid flow phantom. We then measured brain activity during a word-color Stroop task in 15 subjects and obtained brain activation maps. The average signal changes in the channel showing the largest activation were 1.7 × 10^−2^ in ΔOD and 6.6% in CBF.

## Introduction

Cerebral blood flow (CBF) is a critical hemodynamic parameter that reflects the delivery of oxygen and nutrients to brain tissue, supporting neuronal function and brain health. CBF is an important metric of brain health, as abnormal changes in CBF have been associated with numerous pathologies, including ischemic stroke^1,2^, traumatic brain injury^3^, Alzheimer’s disease^4,5^, and Parkinson’s disease^6^. CBF changes also reflect brain function due to neurovascular coupling, a dynamic process where neuronal activity induces localized hemodynamic changes^7,8^. Therefore, non-invasive methods for monitoring CBF can offer great value in both clinical and neuroscience applications. Many methods have been developed to monitor CBF. For instance, magnetic resonance imaging (MRI) based arterial spin labeling is the current state-of-the-art method for CBF mapping. But MRI suffers from low signal-to-noise ratio (SNR), low temporal resolution, and high-cost from the need for high magnetic fields and dedicated, electrically-shielded imaging suites. Similarly, computed tomography (CT) perfusion and positron emission tomography also require expensive scanning hardware and, furthermore, involve ionizing radiation^9^. Transcranial Doppler ultrasound is another method able to infer CBF from velocity measurements in large cerebral arteries, but is also relatively cumbersome, requires expert operators, and does not provide spatial maps of blood flow^10^. Therefore, these existing imaging methods are not preferred for continuous CBF monitoring at the bedside or in naturalistic settings.

Optical methods serve as a promising candidate for CBF monitoring as they enable the development of low-cost, wearable systems that allow for continuous monitoring of the human brain at the bedside or in outdoor environments for naturalistic neuroscience studies. Diffuse correlation spectroscopy (DCS) is an optical method that captures flow information from the temporal dynamics of speckle patterns created by the interference of coherent light that scatters through the brain^11^. In DCS, the temporal speckle intensity autocorrelation function *g*_2_(*τ*) = ⟨*I*(*t*)*I*(*t* + *τ*)⟩/⟨*I*(*t*)⟩^2^ is calculated where ⟨…⟩ indicates an ensemble average, *t* is time, *τ* is the time lag and *I* is the intensity. CBF is directly related to the apparent diffusion coefficient *D*_*B*_ of scatterers that determines the shape of the *g*_2_(*τ*) curve^12,13^. It is non-invasive, convenient to use, and has been applied in numerous clinical studies^14–18^. However, traditional DCS suffers from low SNR, which limits DCS to relatively small source-detector separations (≤25 mm), where the sensitivity to the brain is low, i.e., the signal is more sensitive to the extracerebral (scalp and skull) rather than cerebral blood flow changes. This is because the intensity autocorrelation, *g*_2_(*τ*), decays on the order of approximately 10 microseconds, and traditional DCS utilizes relatively high-cost, high sampling rate, single photon avalanche diodes as detectors to temporally resolve *g*_2_(*τ*), making it expensive for DCS to scale to a multi-channel system capable of covering a large area of the brain, or averaging over multiple speckles to improve SNR. Interferometry has been shown to improve DCS SNR^19,20^, but at the expense of increased system complexity which is not preferred for the development of multi-channel or wearable devices.

Speckle contrast optical spectroscopy (SCOS) is another optical technique that is capable of measuring CBF, which has grown rapidly over the last few years showing competitive advantages over DCS^21–28^. Instead of temporally resolving the intensity autocorrelation function as done in DCS, SCOS calculates the contrast of the speckle pattern over space or time, defined as the ratio between the standard deviation and the mean of the intensities of the speckle images,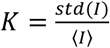, where *std*(…) indicates the standard deviation. Unlike DCS, SCOS measures the integration of the speckle intensity over an exposure time and does not need to temporally resolve *g*_2_(*τ*). Therefore SCOS can achieve higher SNR than DCS (>10x), thanks to the averaging of the contrast over thousands to millions of pixels/speckles available with low-cost complementary metal-oxide semiconductor (CMOS) cameras^29^. Recently, we presented the ability of single-channel SCOS to measure changes in CBF through computational modeling^30^ and experimental validations using a scientific CMOS (sCMOS) camera^25^. We then found that more affordable, commercial CMOS cameras can also be used for SCOS measurements and can achieve accurate measurements^31^. These more affordable detectors make it feasible and accessible to build a multi-channel system to cover a large area with multiple sources and detectors. The resulting overlapping source-detector pairs have photon trajectories that cover different parts of the brain, allowing for spatially specific measurements of CBF changes in the brain.

In this manuscript, we present the development of a multi-channel, high-density SCOS system consisting of 7 sources and 17 detectors that can map CBF or *D*_3_ changes over an area of 7.6 cm by 6.6 cm with a high measurement density (19 mm nearest neighbor source detector separation (SDS), excluding the 8 mm SDS short separation channel). While numerical simulations have suggested the potential of SCOS for tomographic brain imaging^32^, and preliminary studies have demonstrated its ability to measure cerebral blood flow in humans at a single channel^25,33^, mapping of human brain function with high-density SCOS systems has not yet been achieved. We have temporally multiplexed 7 source positions at 14.3 Hz, sufficient to appropriately sample task-induced hemodynamic responses. We demonstrated the spatial specificity of the high-density measurements using a flow phantom that produced a local change in flow velocity. We then conducted human brain function measurements using a word-color Stroop (WCS) task^34^. The results show the capability of our multi-channel, high-density SCOS system to non-invasively map changes in CBF in the human brain, providing a valuable tool for studying hemodynamic responses and brain function in diverse cognitive and clinical applications.

## Results

The schematic of the multi-channel SCOS system consisting of 7 sources and 17 detectors is shown in Fig. 1a. For the source, a galvanometer steered 852 nm laser light to 7 fibers in a multi-mode fiber array projecting to 7 head locations. The distal end of each source fiber was placed in a 3D-printed fiber holder and coupled to a mirror angled at 45° and an acrylic light pipe that directed the light towards the skin (Fig. 1b). The 16 detectors consisted of a bundle of multi-mode fibers that transmitted the speckle patterns formed by reemitted photons on the head. The speckle pattern is imaged onto a camera, with a 4f system to magnify the image (2.2x). For short source-detector separation measurements, the 17th detector used a commercial optode (NIRx Medical Technologies LLC, Berlin, Germany). The optodes were placed in a hexagonal grid with the nearest source-detector separation (SDS), excluding the separate short separation channel (8 mm), at 19 mm, followed by the second nearest SDS at 33 mm (Fig. 1c,d). Similar to fNIRS, the sampling depth is approximately one-half to one-third of the SDS^35,36^; thus for an SDS of 33 mm, the sampling depth is roughly 11-16.5 mm. The short separation NIRx channel was placed adjacent to source 7 at an 8 mm SDS. The diagram of the control system for the multi-channel SCOS set-up is shown in Fig. 1e. More details on the system set-up are provided in the methods section.

**Figure 1.**
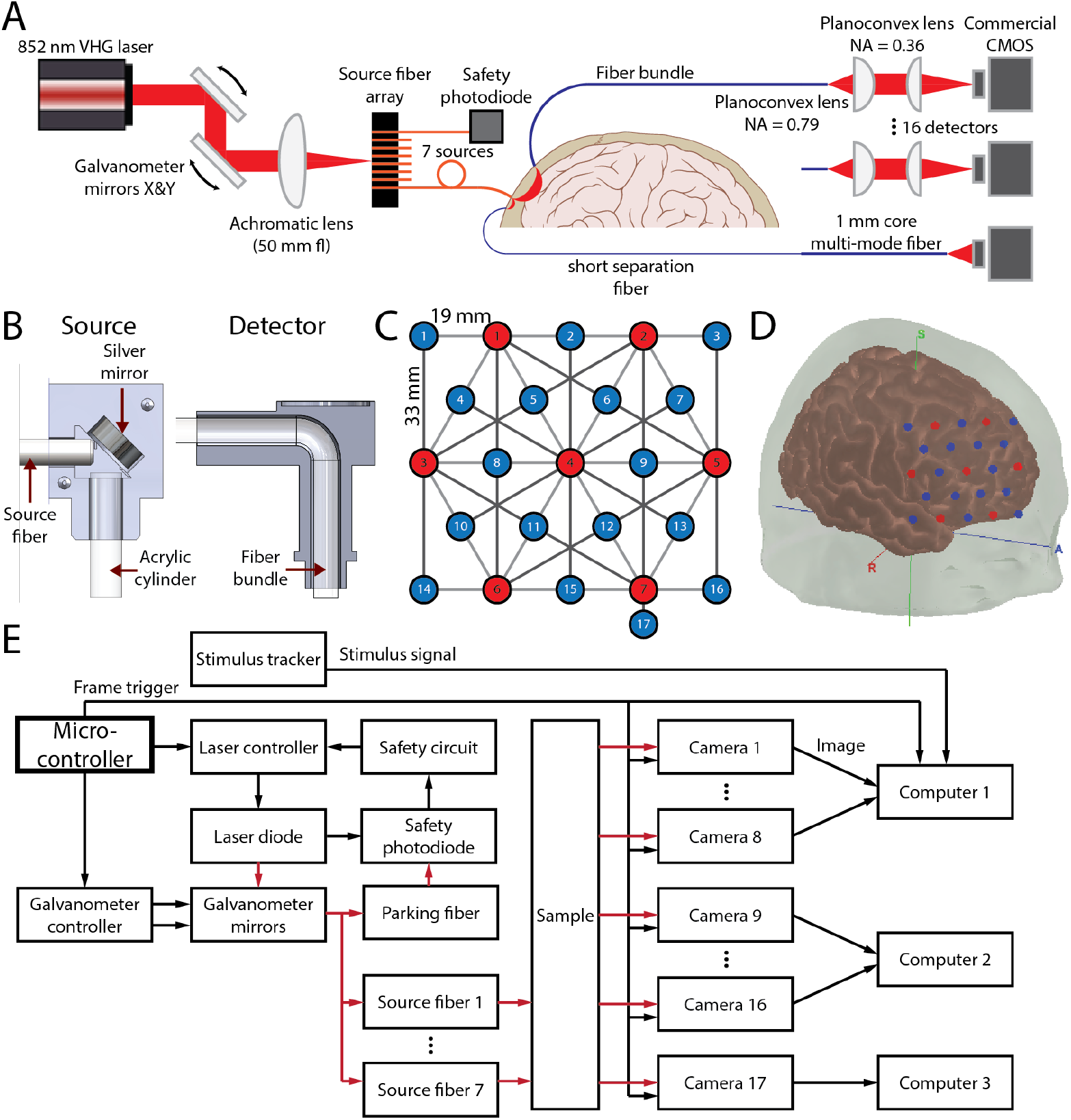
Multi-channel SCOS system setup. (A) Schematic of the multi-channel SCOS system. (B) Cross section CAD diagram of the source fiber holder (left) and detector fiber holder (right). (C) The 76 × 66 mm grid array consisting of 7 sources (red) and 17 detectors (blue). The 17^th^ detector has a SDS of 8 mm from source #7. (D) The location of the source detector grid array on the head for the cognitive task. (E) Information flow diagram of the SCOS control system. Black arrows indicate electrical signals, while red arrows indicate optical signals.

We evaluated the stability of our system’s laser light inputs for the temporal multiplexing scheme utilized to illuminate multiple source positions with a single laser light source. To test the stability during light steering, we used a photodiode to measure the intensity at the distal end of each source fiber while maintaining constant voltage to the galvanometer controllers. Any fluctuations in the time course were attributed to galvanometer instability, shot noise, and/or photodiode read noise. We then repeated the measurement with a trapezoidal waveform galvanometer voltage input instead of a constant voltage. We used a trapezoidal voltage waveform instead of a square waveform to reduce galvanometer mirror vibration caused by large accelerations. This trapezoidal voltage waveform was used to steer the beam between individual source fibers for human measurements as well. We obtained the intensity time course by averaging over 4 ms windows, with each window starting 1 ms after the beginning of the plateau, as shown in Fig. 2a. An example intensity time course from one of the 7 source fibers is shown in Fig. 2a. We aimed to achieve an intensity fluctuation at least 10 times lower than the intensity variations induced by brain activation. The temporal standard deviation of the normalized intensity time course, arising from noise and system instability, increases from 3.3 × 10^−4^ when the galvanometer mirrors remain stationary to 5.6 × 10^−4^ when the galvanometer mirrors move between source fibers. This variation is more than an order of magnitude lower than the expected intensity signal changes of 1 × 10^−2^ to 2 × 10^−2^ during brain activation. In Fig. 2c, we present a baseline phantom measurement using a liquid phantom composed of Intralipid^®^ 20% (Fresenius Kabi USA, LLC, Lake Zurich, IL, USA) diluted to 1.6% v/v in deionized water at room temperature) with a camera exposure time of 4 ms. Fig. 2c shows the average intensity recorded by each detector camera when light was launched sequentially through all the source locations. The intensity level is related to the source-detector separations, showing that the temporal multiplexing scheme is working as expected. For instance, at detector #1, the closest light source is source #1, followed by source #3. As expected, the intensity is highest when source #1 is illuminated and second highest when source #3 is illuminated.

**Figure 2.**
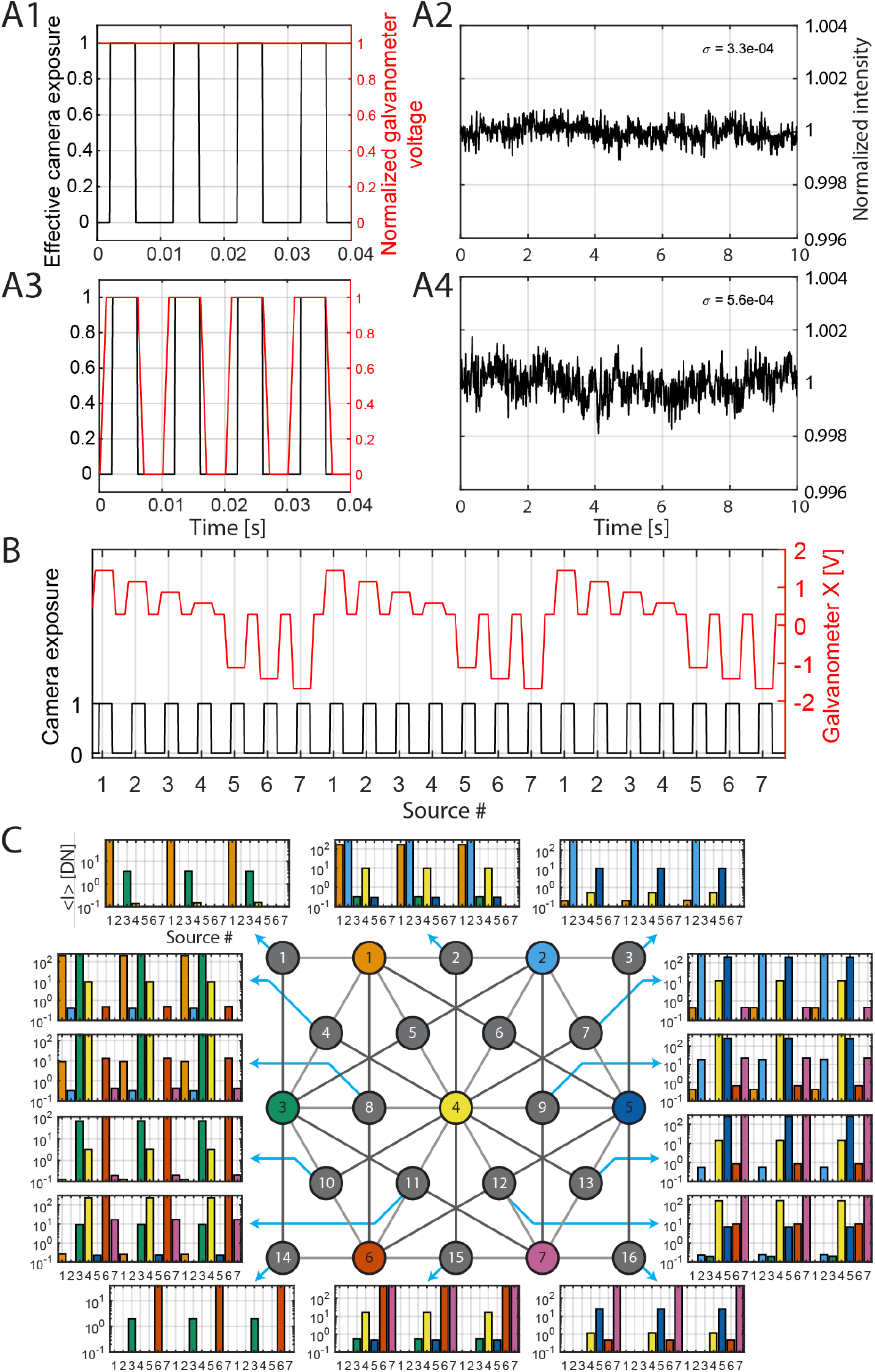
Temporal multiplexing scheme of the sources. (A) Intensity stability with and without galvanometer mirror movement. The time course for the source (red) and detector (black) controls when the galvanometer mirror is not moving (A1) and moving with a trapezoidal form (A3). (A2) and (A4) are the recorded intensity time courses for (A1) and (A2) respectively. (B) Example synchronized camera exposure (1 exposing and 0 not exposing) and galvanometer voltage time courses. The voltage of the galvanometers determines the location of the laser beam relative to the source fiber array, therefore controlling which source fiber is illuminated. (C) The intensity measured at all the 15 detectors using a liquid phantom sample when the 7 sources are illuminated sequentially.

Next, we tested our system’s ability to capture cardiac waveforms from the human head. The source-detector array was placed on the human forehead as shown in Fig. 3a. We show the time courses of ΔOD = − log *I*/*I*_0_, where *I*_0_ is the mean intensity over the measurement period, and fractional change in blood flow,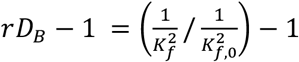, where 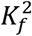 is the fundamental contrast squared, 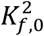 is the fundamental contrast squared at baseline, and *rD*_*B*_ is the relative blood flow index, for the channels with SDS of 33 mm in Fig. 3b. We can clearly visualize the cardiac waveforms simultaneously in all the channels for both ΔOD and *rD*_*B*_ − 1.

**Figure 3.**
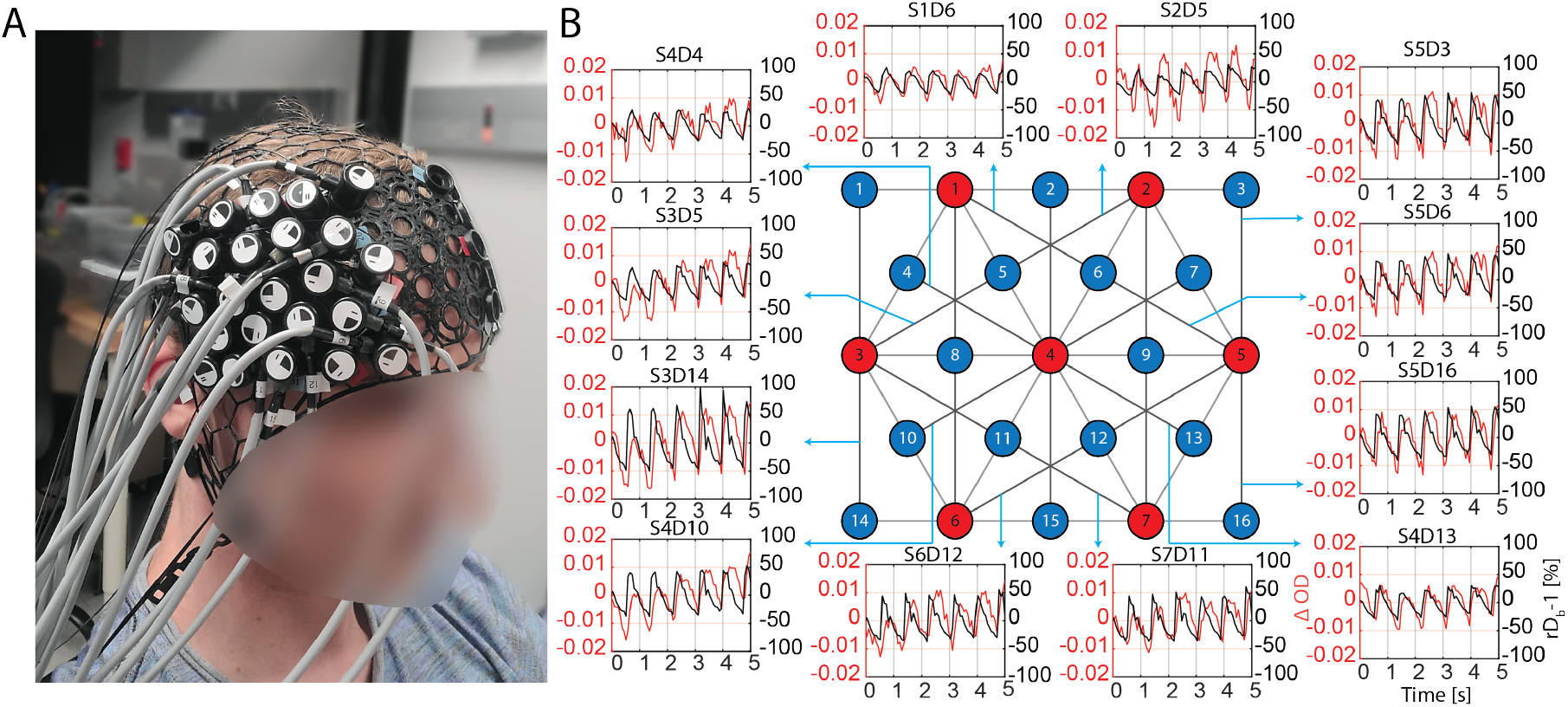
Cardiac waveforms measured in 33 mm SDS channels. (A) A participant wearing the multi-channel SCOS system on the right forehead. (B) Cardiac signal in ΔOD (red) and rD_b_-1 (black) shown from multiple 33 mm SDS channels. The measurement rate is 14.3 Hz.

To demonstrate our system’s capability to localize flow variations, we used a dynamic phantom sample (same as that used in Fig. 2) placed in an acrylic tank with a transparent, thin-walled tube (0.082 inches inner diameter) positioned 12 mm from the tube center to the fiber tips of the high-density probe. The tube was subjected to bulk flow at 2 cm/s for 30 s, followed by 30 seconds of no flow to allow baseline recovery. This cycle was repeated 6 times, with the time course of the flow shown in Fig. 4a. The time courses of the changes in *rD*_*B*_ − 1 and ΔOD from -10 to 45 seconds (with 0 second indicating the onset of the flow) are shown in Fig. 4c and d, respectively. These averages, taken over 6 blocks, are displayed in channel space for SDSs of 19 mm and 33 mm, and for the orientations of the source-detector layout relative to the tube (0^0^, 45^0^, 90^0^) as illustrated in Fig. 4b. The profiles were obtained using the PlotProbe2 function in the open-source data processing software Homer3^37^. We observe that the *rD*_*B*_ − 1 increases from 0 to 30 seconds for channels (i.e. source-detector pairs) overlying the tube, while no change is seen in ΔOD, as expected, since there were no absorption changes. This illustrates that our multi-channel, high-density SCOS system can measure spatiotemporal *rD*_*B*_ − 1 changes, which is an independent parameter not accessible with functional near infrared spectroscopy (fNIRS) that only measures ΔOD which is related to blood volume or hemoglobin concentration changes. Additionally, we see that the amplitude of “activation” across channels varies with different orientations of the source-detector layout, with channels spatially closer to the tubes consistently showing larger *rD*_*B*_ − 1 increase. This highlights the capability of the multi-channel SCOS system to map spatially inhomogeneous dynamic changes.

**Figure 4.**
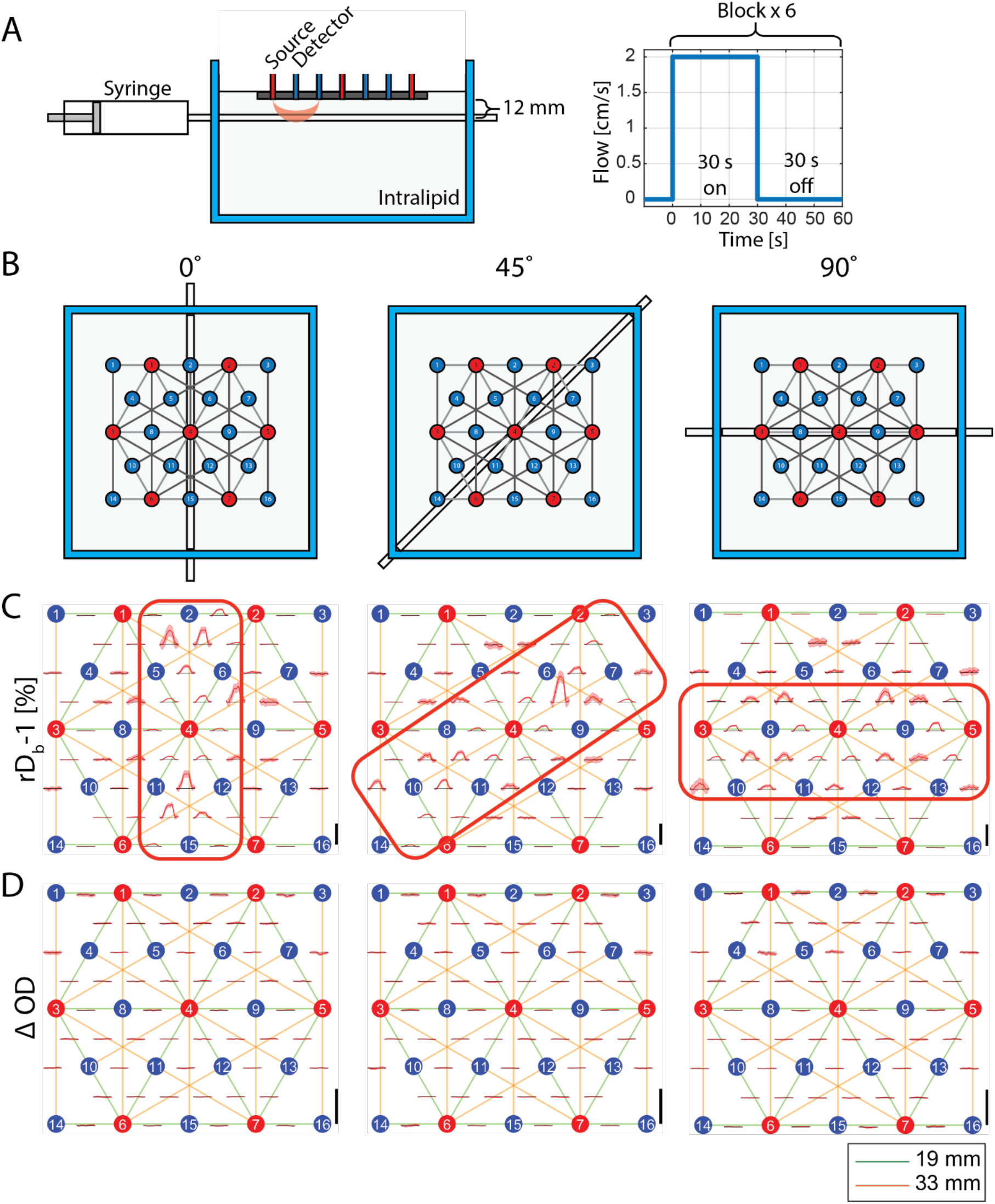
Demonstration of the measurement of flow changes using the multi-channel SCOS with a dynamic phantom sample. (A) Dynamic phantom flow measurement setup, and the time courses of the flow within the tube. (B) The orientations of the source-detector layout relative to the tube at 0^0^ (left), 45^0^(middle), and 90^0^ (right). (C)-(D) rD_b_-1 (C) and ΔOD (D) Time course signals averaged over 6 blocks for the 19 mm and 33 mm SDS channels, shown in the PlotProbe2 profile using the Homer 3 software. Shaded area is the standard error of the time course over the blocks. The red rectangles highlight channels where changes of the signals are expected. The y-scale shown as a black vertical bar on the right lower corner of each plot represents 20% for *rD*_*B*_ − 1 and 0.02 for ΔOD, respectively.

Finally, we conducted human brain function measurements using a WCS task (Figs. 5a,b) consisting of a congruent (easy) and an incongruent (hard) task condition, as described in the Methods section. In congruent trials, the participant chooses the word with matching font color and meaning. In incongruent trials, the participant chooses the word in the top row with meaning matching the font color of the word in the bottom row. We chose to use WCS task for its ability to induce frontal lobe activation and provide feedback on the participant’s engagement through their answer accuracy. Out of the 19 participants measured, data from 15 participants, which had more than half of the blocks not contaminated by motion artifacts, were further utilized for analysis. The participants overall showed high task performance, with average answer accuracy of 96.6% (standard deviation of 4.9%, range from 82% to 100%). When divided into congruent and incongruent task performance, participants chose the correct answer 99.5% (standard deviation = 1.1%) for congruent tasks and 93.9% (standard deviation = 9.5%) for incongruent tasks. The collected speckle images were preprocessed to obtain mean and variance for each 7 by 7 window, then the preprocessed data were analyzed to obtain both ΔOD and *rD*_*B*_ − 1. We linearly regressed out the short separation channel from both signals, thereby obtaining ΔOD and *rD*_*B*_ − 1 with reduced scalp signal contaminations. *rD*_*B*_ − 1 was then low pass filtered (0.2 Hz) and averaged over blocks and participants to obtain activation time courses over channel space for 15 participants (Figs. 5 c,d). Both the channel space maps (Figs. 5 c,d) and reconstructed images (Figs. 5 f,g,i,j) show group-averaged results across the 15 participants. For each signal type (ΔOD, *rD*_*B*_ − 1), significant channels were selected as those with mean activation at t > 10s and t < 15s greater than 1.96 times standard error corresponding to 95% confidence level. The spatial distribution of the significant channels for ΔOD covered both lateral and medial regions, while the significant channels for *rD*_*B*_ − 1 covered only the lateral regions, with six 33 mm SDS channels overlapping between the two signal types. The average ΔOD activation of those channels under the incongruent task was (4.2 ± 1.8) × 10^−3^ (*p* = 0.033, n = 15) for 19 mm SDS and (8.0 ± 3.1) × 10^−3^ (*p* = 0.021, n = 15) for 33 mm SDS while the average *rD*_*B*_ − 1 activation of those channels under the incongruent task was 3.3 ± 1.2% (*p* = 0.014 n = 15) for 19 mm SDS and 3.5 ± 1.3% (*p* = 0.013, n = 15) for 33 mm SDS, with results shown as (mean ± standard error). Therefore, there is significant brain activation measured for both 19 mm and 33 mm channels considering the significance level of 0.05. The same channels did not have significant activation under the congruent task with ΔOD changes of (2.5 ± 1.9) × 10^−3^ (*p* = 0.206, n = 15) for 19 mm SDS and (3.6 ± 2.5) × 10^−3^ (*p* = 0.179, n = 15) for 33 mm SDS, and *rD*_*B*_ − 1 changes of those channels were 0.4 ± 0.7% (*p* = 0.572 n = 15) for 19 mm SDS and 0.5 ± 0.8% (*p* = 0.531, n = 15) for 33 mm SDS. The signal changes for the channel with the largest activation for each subject are listed in Supplemental Table 1 for the incongruent task. The subject averaged signal changes in the channel showing the largest activation were 1.7 × 10^−2^ in ΔOD and 6.6% in r*D*_*b*_ − 1.

**Figure 5.**
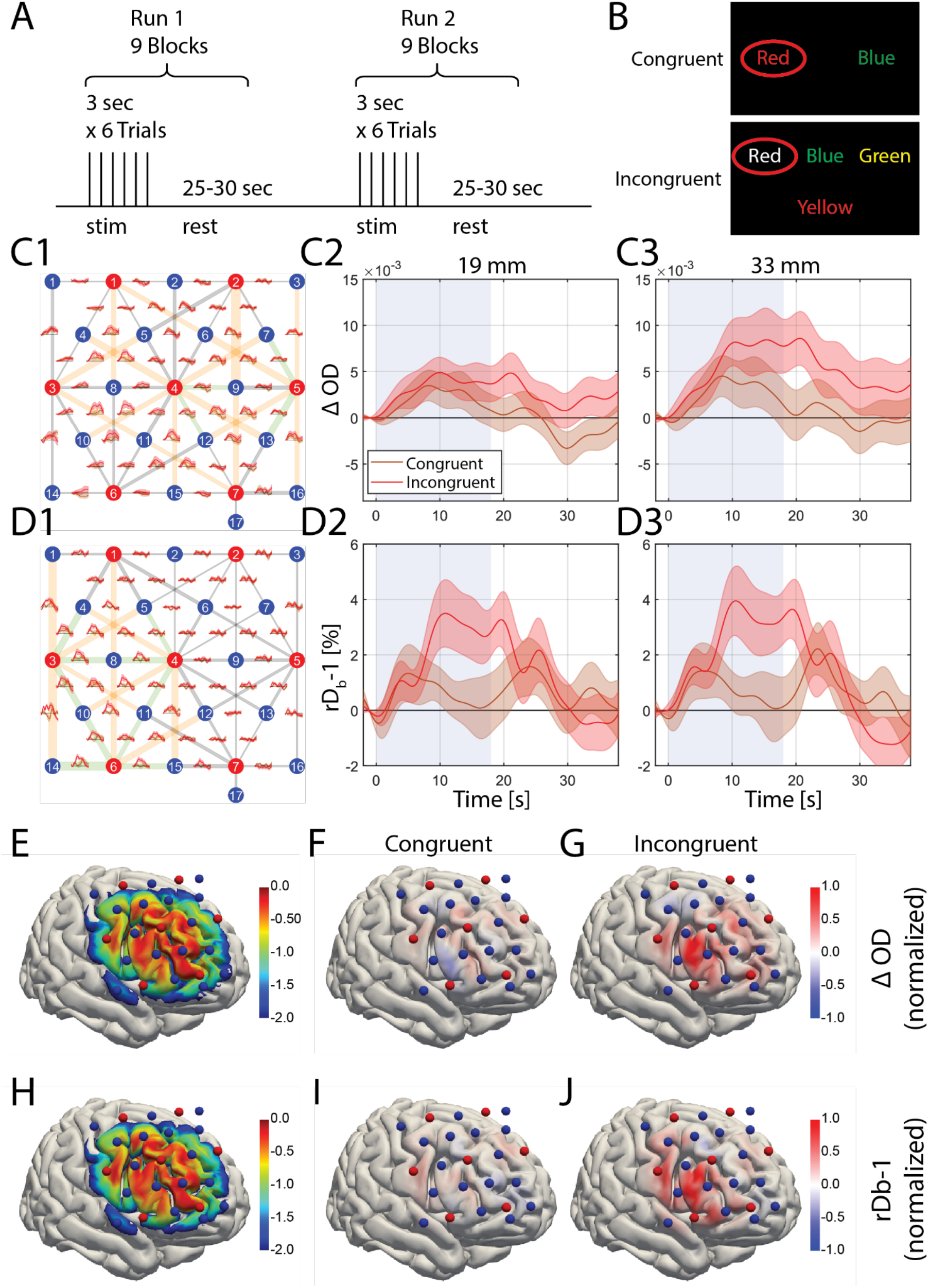
Task based human brain activation measurement using the multi-channel SCOS system. (A) Illustration of the task. Each participant has 2 runs consisting of 9 blocks each, and each block is composed of 6 WCS task trials each lasting 3 seconds, followed by a rest time of 25-30 seconds. (B) Demonstration of the congruent (upper) and incongruent (lower) WCS task. (C)-Group-averaged channel space (C1) ΔOD and (D1) *rD*_*B*_-1 source-detector channel space map, with 19 mm SDS channels (green) and 33 mm SDS channels (orange) demonstrating channels with significant ΔOD activation (mean > 1.96 standard error) highlighted. The shaded area indicates standard error across subjects. The average results over these highlighted channels are shown in (C2) 19 mm SDS (C3) 33 mm SDS for ΔOD, and (D2) 19 mm SDS (D3) 33 mm SDS for *rD*_*B*_-1. (E) Normalized sensitivity map for ΔOD, normalized, log_10_ scale. (F,G) Group-averaged WCS induced ΔOD (t > 10s & t < 15s) reconstructed image for congruent (F) and incongruent (G) tasks. (H) Normalized sensitivity map for *rD*_*B*_-1, normalized, log_10_ scale. (I,J) Group-averaged WCS induced *rD*_*B*_-1 (t > 10s & t < 15s) reconstructed image for congruent (I) and incongruent (J) tasks.

We utilized a Monte Carlo model integrated with Cedalion^38^, which is an open-source software for fNIRS data processing, to reconstruct both the ΔOD and *rD*_*B*_ − 1 images (Figs. 5 e-j). The sensitivity matrices for ΔOD (Fig. 5 e) and *rD*_*B*_ − 1 images (Fig 5 h) appear similar. This is expected, as although the signals originate from different tissue properties, they are derived from the same photons measured within a given channel. Therefore, spatial sensitivity is unlikely to significantly differ. Additionally, both sensitivity matrices show a lack of sensitivity at the edges of the array, which is expected, as there are fewer overlapping channels at the border. In the reconstructed images, the spatial distribution of *rD*_*B*_ − 1 (Fig. 5 j) showed more focal activation patterns on the lateral side, while ΔOD (Fig. 5 g) exhibited a slightly broader, more diffuse pattern of activation. The spatial distribution of the activation observed in channel space is consistent with the reconstructed image, both showing lateral activation. This activation is roughly localized to the dorsolateral prefrontal cortex (DLPFC).

## Discussion

Our multi-channel SCOS system demonstrates capabilities for CBF monitoring with high spatial specificity and temporal resolution, marking an advancement in the field. We expanded upon our prior work with a one-source, one-detector SCOS system by using a galvanometer to illuminate 7 source fibers and 17 CMOS cameras as detectors. We demonstrated the multi-channel SCOS system’s ability to measure physiological signals, as well as its spatial specificity to underlying flow change with a dynamic phantom experiment. The human applicability of the system was presented by non-invasively measuring a WCS task-induced spatial activation pattern of CBF changes in humans.

For the liquid phantom experiment, we measured underlying flow with the multi-channel SCOS system at different tube orientations (0°, 45°, 90°) and showed that channels closer to the tube have higher *rD*_*B*_ − 1, in line with the expectation. For example, channels that do not overlap with the tube spatially at 0° but do at 90° (S3D5, S3D11) show *rD*_*B*_ − 1 increase only with the tube at 90° orientation. The magnitude of the *rD*_*B*_ − 1 increase was, in general, larger for 33 mm SDS channels than 19 mm SDS channels, consistent with the higher sensitivity of the larger SDS channels to deeper flow change. One limitation of this experiment is that flow through the tube primarily increases convective flow, rather than diffusive flow. Meanwhile, prior Monte Carlo simulation of DCS showed primary contribution from diffusive flow in biological tissue^39^, meaning there is a mismatch in the cause of the *rD*_*B*_ − 1 increase between phantom and *in vivo* measurement. Another limitation is that the phantom is homogenous in both scattering coefficient and flow speed at baseline, which is not anatomically representative of the head, which has different scattering coefficients and flow speeds for the scalp, skull, and brain. Lastly, vibration or motion of the tube during the flow condition may result in fluctuations in ΔOD and r*D*_*B*_-1. However, we expect any change in speckle dynamics from motion of the tube or vibration of the pump would be much smaller compared to the 2 cm/s flow through the tube^7^.

Prior human studies measuring cognitively-activated cerebral blood flow change used visual stimuli and motor task. We chose WCS task because it activates the prefrontal cortex which is in a hair-sparse region of the head, allowing for easier optode placement without compromising photon flux, and because participant engagement can be measured through their task accuracy. In the image reconstruction result, activation of the DLPFC was observed. This activation is consistent with fNIRS measurements that also showed DLPFC activation during the WCS task^34,40,41^.

Our human measurements show statistically significant channels distributed differently for ΔOD and *rD*_*B*_ − 1. While significant ΔOD channels are present both medially and laterally, significant *rD*_*B*_ − 1 channels are only located laterally. The same pattern is seen in the reconstructed image, where the ΔOD map shows a broader activation pattern than *rD*_*B*_ − 1. One possible explanation for the difference in the spatial distributions of ΔOD and *rD*_*B*_ − 1 is that measurements were conducted at a single wavelength (852 nm) and ΔOD alone cannot accurately map brain activation regions. Addressing this limitation requires the integration of SCOS and fNIRS with two or more wavelengths to simultaneously map *rD*_*B*_ − 1, oxyhemoglobin (HbO), deoxyhemoglobin (HbR), and total hemoglobin (HbT) changes. Additionally, the regularization needed for image reconstruction could be different between ΔOD and *rD*_*B*_ − 1. These need to be validated in the future with more subject measurements and comparing SCOS with fNIRS. In this work, we have ignored the potential variation of the speckle contrast and the sensitivity matrix induced by changes of absorption during brain activation^42^. This is a higher-order effect that non-linear image reconstruction methods can handle. However, for the small absorption changes of a few percent that occur during brain activation, this will generally produce negligible changes in the spatial sensitivity profile and can thus be ignored. Future work could explore the limits when this assumption would break down.

Several technical considerations and limitations merit discussion. One practical challenge is the variation in coupling efficiency between source fibers, leading to differences in light intensity among the seven sources (approximately 20%). While this variation affects the signal-to-noise ratio across different channels, it does not impact the mean values of OD and *rD*_*B*_ − 1. This is because both the numerator and denominator of *e*^−Δ56^ are linearly proportional to the baseline intensity, and *rD*_*B*_ − 1 remains input light intensity-independent if noise correction is properly applied.

The temporal resolution of our system is currently constrained by the multiplexing strategy employed. Each source position requires 10 ms using trapezoidal waveforms to control the galvanometer mirrors, resulting in a frame rate of 14.3 Hz to cycle through all seven sources. Although this resolution is sufficient for capturing the hemodynamic responses during brain activation relevant to our study, future systems with more channels may need multiple laser sources operating in parallel to maintain adequate temporal resolution for other applications. Regarding spatial resolution, our current setup covers a 7.6 cm x 6.6 cm area with source-detector separations (SDS) ranging from 19 mm to 33 mm. The spatial resolution of this multi-channel SCOS system is expected to be comparable to that of fNIRS with a similar optode density^43^, though coarser than traditional neuroimaging methods like fMRI. This limitation arises from the number of channels that cover different parts of the brain. Increasing the optode density in the future could improve the spatial resolution^44^.

Motion sensitivity is another challenge, as observed in most optical imaging techniques. Head motion affects the optode-to-head coupling, resulting in signal artifacts^45^. While we minimized head motion during our human participant measurements using a chin mount and instructions to the participants, we still observed substantial intensity fluctuations in more than half of the blocks in 4 of the 19 participants, leading to their data being excluded from group analysis. A potential solution could involve a fiber-free SCOS system that is fixed on the head to reduce motion artifacts, which also allows for more naturalistic behavior and expanding the scope of neuroscience and clinical applications^33^.

Furthermore, the current implementation requires careful positioning of the optodes to ensure good coupling with the scalp and proper threading through the hair. Although we have optimized the fiber holders and mounting system with spring tops, maintaining consistent coupling during long measurement sessions remains a practical challenge that needs to be addressed in experimental design.

The SCOS system uses 17 cameras, each with 2 million pixels and 16 bits per pixel, capturing data at 100 frames per second over 14 minutes. This setup generates approximately 12 terabytes (TB) of data per participant. Measurement duration is therefore limited by the available computer storage. Our preprocessing pipeline, which calculates the windowed mean and variance for each image, reduces the data size to hundreds of gigabytes. However, this process is currently implemented post hoc. A real-time implementation of preprocessing would enable significantly longer measurement time; however, extensive validation is needed to ensure that the proper noise correction is maintained during real-time image processing. Data analysis methods commonly used for fNIRS, such as principal component analysis^46^ and general linear models^47,48^, may also be applicable to SCOS data analysis^49^, which will be tested in the future.

Another confounding factor is the potential for participants to develop increasing familiarity within the task over the course of the experiment, which may influence the hemodynamic responses. Additionally, participants often reported feeling increasingly tired as the experiment progresses, which could further impact the measurements. To characterize the potential confounds, the hemodynamic response was compared between the first and second run of the experiment (Supplemental Figure 2). For the congruent task, larger activation was observed in run 1 ((6.2 × 10^−3^ for ΔOD and 2.2% for *rD*_B_ − 1) than in run 2 with (1.7 × 10^−3^ for ΔOD, −0.7% for *rD*_*B*_ − 1), suggesting that repeated exposure could made the task insufficiently challenging to induce activation, especially for relatively easy tasks.

Looking ahead, this multi-channel SCOS system shows potential for advancing functional neuroimaging. Future work includes implementing real-time preprocessing algorithms to extend measurement durations and improve data management efficiently. Future advancements may focus on designing optodes that can better penetrate through the hair, as well as enhancing the SNR to ensure sufficient photon detection despite absorption by the hair. Adding more channels for higher density could improve spatial resolution, while more coverage would allow for whole head measurements. Potential future research applications include mapping of cerebral activation in response to various stimuli (finger tapping, visual tasks), and comparing cerebral blood flow and cerebral blood volume based resting-state functional connectivity^50^. With further development, potential clinical research applications could include monitoring of cerebral hemodynamics under conditions of disrupted cerebral homeostasis, such as during anesthesia, acute neurological disorders like cerebral infarction, and chronic neurological disorders such as Parkinson’s disease and Alzheimer’s diseases^4,6^.

## Methods

### Multi-channel fiber-based SCOS system

The schematic of our multi-channel fiber-based SCOS system is illustrated in Fig. 1. The input laser light source (Thorlabs free-space VHG, 852 nm) is first collimated using a planoconvex lens (C110TMD-B Numerical Aperture NA = 0.4), then passed through an optical isolator (IO-5-850-VLP). The light then enters a galvanometer scanner (GVSK2-US) and is focused down using a planoconvex lens (AC254-050-B NA = 0.25). The galvanometer couples the focused light into a multi-fiber termination push-on (MTP) fiber array containing 12 multimode fibers (62.5 μm core diameter, 0.22 NA), 7 of which are coupled to acrylic light pipes of 3 mm diameter that deliver the light to a sample such as the human head. We utilized custom-made fiber bundles (∼3770 strands of 37 µm core diameter multimode fiber, 0.66 NA) with circular shape on the proximal end (∼2.5 mm diameter) to facilitate scalp contact through the hair and rectangular shape on the distal end (∼1.64 × 3 mm) to match the rectangular shape of the CMOS sensors used as detectors. We imaged 16 bundles from different locations on the human scalp onto 16 CMOS (Basler a2A1920-160umPRO) cameras. The fiber bundle was imaged using a 4f system (ACL25416U-B NA = 0.79, 38-421 NA = 0.36) as shown in Fig 1a. The cameras operate at 10-bit resolution with 16 dB gain and a black level of 25 analog-to-digital unit (ADU) with an average read noise of 2.1 e-across the sensor. The plano-convex lens closer to the fiber bundle was moved slightly off-focus (2-3 mm) to defocus the imaged speckle to fill the gaps between individual fibers within a single fiber bundle, as shown in Supplemental Figure 1, resulting in a reduction in spatial heterogeneity of the illumination on the sensor. The s/p and *β* for the system have been calibrated from 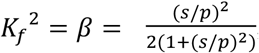, where *β* is the system coherence parameter, for unpolarized light obtained using a static phantom sample at high photon flux values to reduce errors introduced by noise^30^. The values of s/p are measured to be ∼0.69, and *β* to be ∼0.16 for all the cameras. The sources and detectors are arranged in a symmetrical hexagonal grid as shown in Fig. 1c. Additionally, a short separation channel is included using a commercial short-separation fiber (NIRx Medical Technologies LLC, Berlin, Germany) holder module (50-4328 R3) mounted around the source optode 7 to achieve an SDS of 8 mm for short separation regression^51^. The short-separation fiber holder module attaches directly to the source grommet on the cap creating an 8 mm SDS channel that is primarily sensitive to the scalp, making it ideal for short separation regression. The NIRx module contains a fiber bundle that is coupled to a 1 mm diameter multimode subminiature version A (SMA) optical fiber (0.22 NA) that projects light directly onto a camera. This fiber was then connected to the camera sensor, positioned 15 mm away from the fiber tip. Due to the smaller SDS, the amount of light in the short-separation channel is significantly higher, allowing for direct projection onto a 17^th^ CMOS camera without requiring the optimization of s/p.

The source focal spot size was measured by reducing the laser diode current to 3.2 mA and placing the camera at the same z-plane as the fiber array. Using a 6.2 mm focal length collimation lens and a 50 mm focal length achromatic focusing lens achieved a focal spot of 55 µm full-width at half-maximum (Supplemental Figure 1). The laser diode had a maximum output power of 600 mW. The power achieved at the source fiber holder was approximately 270 mW. This reduction is attributed to ∼80% transmission efficiency through the isolator, ∼70% fiber coupling efficiency, and ∼80% mirror and acrylic coupling efficiency. To meet with the American National Standards Institute (ANSI) safety standards, each fiber is illuminated for 5.2 ms every 70 ms, resulting in a 7.4% duty cycle, which reduce the average power to 20 mW. This ensures both the peak and average power are within the ANSI safety limits.

The 7 source fibers and 17 detector fiber bundles are placed in a 76 mm by 66 mm rectangular grid (Fig 1c), resulting in 28 pairs of 19-mm SDS channels and 22 pairs of 33-mm SDS channels. For human measurements, the grid is placed on the right forehead to target the right lateral frontal lobe (Fig 1d).

### Multiplexing scheme using the galvanometer mirrors

Multiplexing of the sources is achieved using a galvanometer and a fiber array. As shown in Figure 1, the laser source is collimated then sent into the galvanometer which is controlled by an Arduino connected to an external digital to analog converter (DAC) using a custom designed printed circuit board (PCB). The two galvanometer mirrors, each rotating orthogonal to the other to achieve two-dimensional coverage, steers the light into one of the fibers, where a focusing lens focuses the collimated beam to a spot for improved coupling with the fibers. The beam is sequentially steered to sources 1 through 7, pausing at a safety fiber between each source (details on the safety fiber’s purpose are provided below). Each fiber is routed to a unique location on the head, enabling multichannel measurements through temporal multiplexing.

To achieve optimal coupling, a galvanometer optimization protocol is conducted to locate the coordinates for each fiber. This is repeated before each measurement. The protocol is controlled by a home-made Arduino script. The Arduino receives input from a Thorlabs power meter (Thorlabs PM400), recording from a photodiode (Thorlabs S121C) at which the source fiber is pointed to measure the amount of light from a given source fiber. First, the estimated coordinates are input to the Arduino. Then the galvanometer steers the beam in a spiral radiating outward from the starting coordinate until the threshold of approximately 1 mW is reached. Once the threshold is reached, the galvanometer steers the beam in a line approximately 20 µm in both directions from the starting coordinate. The X coordinate is then set to the peak value along the line scan. The process is repeated for the Y direction. This is repeated until there is no coordinate with higher intensity in either X or Y. Finally, for fine tuning, a square grid of about 40 by 40 µm centered at the highest intensity coordinate is generated, and the laser is scanned throughout the grid. The peak value in the grid is selected as the final coordinate for the source fiber. This process is repeated for all source fibers and the safety fiber, which takes approximately 10 minutes. The coordinates are then saved for later use for the measurements.

### Time synchronization

Because brain activation is measured as an average of the blocks in the task, it is important to synchronize the frames collected by the camera with the light source illumination and the onset of the WCS tasks. This is done by using the Arduino, connected to the PCB, as the primary device that sends trigger signals to the cameras, the galvanometer, and the Data Acquisition (DAQ) device. A stimulus tracker (StimTracker Quad, AD Instruments) is connected to a photodiode that tracks the brightness of a corner of the computer screen that the participant uses for the tasks. The corner of the screen is filled with a white rectangle when the stimulus is given and a black rectangle when there is no stimulus, providing the stimulus tracker with the onset of the stimulus. The DAQ device receives both the transistor-transistor logic signal from the stimulus tracker and Arduino’s trigger signals. This provides the camera frame corresponding to the onset of each stimulus.

### Pulsing strategy to improve photon flux

We utilized a laser pulsing strategy to improve photon flux while staying within the ANSI laser tissue exposure limits. Before measurements, the galvanometer directs the laser onto an eighth fiber connected to a photodiode that controls a safety circuit. During measurements, every 10 ms, the galvanometer steers the light to this eighth fiber for 5.2 ms. If light does not reach the eighth fiber within the 10 ms interval, the safety circuit shuts off the laser to prevent illuminating the human tissue for an excessive duration. Additionally, the temporal multiplexing ensures that a different source is illuminated during each 10 ms period, with the same source only being illuminated again after all the 7 sources have been cycled through, thus achieving a duty cycle of 5.2 ms/ (7× 10 ms)=7.4% for each source. This enables a higher peak power to be sent to the head during acquisition while keeping a lower safe average power and ensures that both the average and peak power are within the ANSI safety limit. Looping through the 7 sources sequentially reduces the measurement rate from 100 Hz to 14.3 Hz for acquiring measurements from all sources.

### Acquisition software

Camera frames are acquired using a home-made Python script employing the PyPylon module (a Python library). The script performs parallel processing, acquiring frames from each camera simultaneously. It also performs multicore processing assigning 2 central processing unit (CPU) cores to each camera for recording. For each camera, the script collects frames and stores them in random access memory (RAM) until 1000 frames are accumulated, at which point they are saved simultaneously. The number of frames collected before saving is a trade-off: collecting too many frames risks exhausting the RAM and losing new incoming frames, while collecting too few frames results in frequent saving, which can delay processing and cause a buildup of frames in RAM, also leading to data loss. As such, a buffer size of 1000 frames was chosen from experience. We found that the required data transfer rate could be sustained by saving data from two cameras to a single hard drive. Each computer utilized 4 hard drives to handle data from 8 cameras simultaneously. This setup required three computers: two for collecting data from the 16 cameras in the high-density patch, and a 3rd computer for managing the short separation camera and running the WCS cognitive task. At the end of each run, a timestamp file was saved for all the cameras, recording the acquisition time for each frame. This file is used during data analysis to identify and correct any potential lost frames.

### Data analysis pipeline for multichannel fiber-based SCOS

Because of the large data size of the raw speckle images (approximately 12 TB per 7 min measurement), data analysis requires substantial computational resources. We first pre-process the data on the computers used for measurements to reduce data size, and then we can transfer the data to other computers to calculate OD and *rD*_*B*_ time traces.

During preprocessing, the timestamp of each frame is first used to correct for any missed frames. Next, the temporal mean dark image, obtained by recording frames with the laser turned off after each measurement, is subtracted from each camera frame to account for the contribution of dark currents. The windowed temporal variance of the dark images and mean images calculated as the temporal mean of all frames for a given source illumination are also saved for noise correction. Finally, pixels are grouped into 7×7 windows, and the mean and variance of each window are calculated and stored. This reduces the total data size by roughly 24-fold.

To perform proper noise correction, the contrast squared from shot noise, read noise, quantization noise, and spatial heterogeneity are calculated so they can be subtracted for each window as shown below in Eqs. 1-6^25,30^.

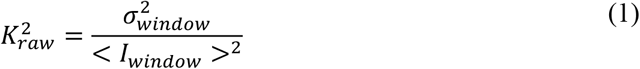

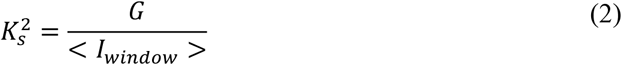

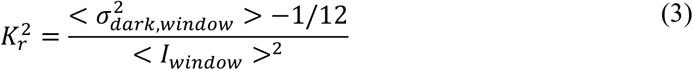

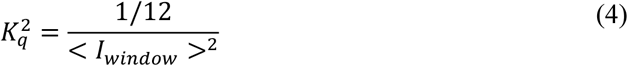

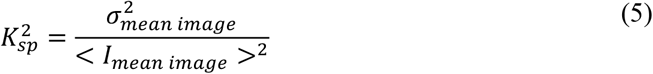

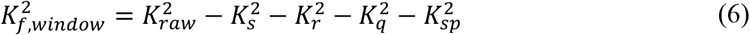

Where 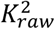 is the contrast squared from the raw dark subtracted data before noise correction, 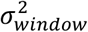 is the windowed variance, < *I*_*window*_ > is the mean intensity of the window, 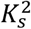 is contrast squared from shot noise, *G* is the camera gain, 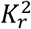 is the contrast squared from read noise, 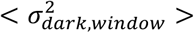 is the spatial mean of the temporal variance of the dark image within a window, 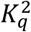 is the contrast squared from quantization noise, 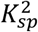is the contrast squared from spatial heterogeneity, 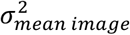 is the spatial variance of the temporal mean image, < *I*_*mean image*_ > is the spatial mean intensity of the temporal mean image, and 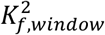 is the fundamental contrast squared due to dynamics in the scattering medium for each 7 by 7 window, which is of interest. The contrast terms are weighted by the mean intensity of the window squared, ⟨*I* _*window*_ ⟩^2^, and normalized to provide higher weight to windows that receive more photons as shown below in Eqs. 7 and 8.

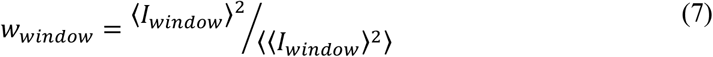

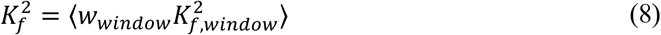

The fractional change of blood flow *rD*_*B*_ − 1 is calculated as the reciprocal of the contrast squared, normalized by the baseline, minus one (Eq. 9).

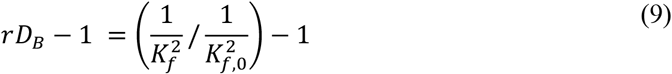

Where 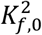 is the temporal mean of 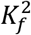 averaged over a baseline time interval before activation.

Motion artifacts result in high-amplitude high-frequency spikes in the optical data, which contaminate results. They are readily distinguishable from physiological fluctuations, which are much lower in both amplitude and frequency. As such, blocks containing motion artifacts were manually identified and removed. Subjects with over 50% of blocks contaminated by motion artifacts were excluded from further analysis, resulting in 15 subjects included out of 19 measured. For the remaining 15 subjects, 83.3% of the blocks (225 out of 270 total blocks) remained after pruning motion artifacts. At low photon counts, besides the low SNR, there is a larger error in the calculation of 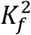 due to the nonlinearity in the photon transfer curve^31^. For this reason, source-detector pairs with mean camera counts lower than 4 digital units were excluded from the analysis, resulting in 96.1% (1442 out of 1500 total source-detector pairs for the 15 subjects with 2 runs each) usable source-detector pairs.

### Camera characterization

Camera characterization is done to plot the photon transfer curve that enables the accurate subtraction of shot noise and read noise contributions from 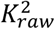^31^. A 785 nm light-emitting diode (LED) is mounted to one port of an integration sphere, with a camera connected to the other port. To limit the range of angles of light incident on the detector surface, the camera is attached to a long lens tube (∼30 cm in length). The LED is connected to a variable power supply, and the voltage to the LED is increased incrementally such that the light intensity recorded from the camera spans the full-scale range of the camera. In this way, the camera intensity variance for each intensity level can be quantified, thus constructing the photon transfer curve.

The entire process is automated using a home-made python script. The script defines a logarithmic scale of intensity values that span the full dynamic range of the camera. The script is responsible for both controlling the power supply and acquiring camera frames. It first acquires dark frames with zero voltage supplied to the LED to measure the dark noise. Then it increments the LED voltage by 1 mV. The intensity is then averaged across the entire frame, and, if the intensity threshold defined by the logarithmic scale is met, the software collects camera frames at that intensity. A logarithmic range is used because typical human measurements occur in the low photon count range, thus given a finite number of measurement points, it is more valuable to have a denser coverage at lower counts. Camera frames are acquired using the PyPylon module, and the power supply is controlled using the PySerial module (a python library).

### Static phantom s/p measurement

Static phantom measurements were conducted to measure the s/p ratio for each camera. A static phantom mimicking human tissue optical properties (absorption coefficient µ_a_ = 0.0188 *mm*^−1^, reduced scattering coefficient µ_s_^’^ = 0.0736 *mm*^−1^) was clamped to the optical table. Next, one of the fiber connector physical contact (FCPC) ends of the source fiber bundle was clamped to the phantom. A detector optode containing the detector fiber bundle was placed 19 mm away from the source fiber. The setup was tightened meticulously to ensure minimal motion to prevent decorrelation of the speckle pattern within the duration of the exposure time. This was verified by monitoring the speckle pattern in real time and ensuring that any observed fluctuations were occurring on time scales slower than a few seconds. Once this was verified, 100 camera frames were recorded with an exposure time of 20 *μs* using a current of 983 mA. To obtain s/p, first 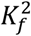 is calculated as described above in Eq. 1-6, except 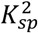 is not subtracted, since for static phantom the temporal mean image is the same as the speckle pattern itself thus it does not provide an estimation of the non-homogeneity in the illumination. We have found that this contribution is three orders of magnitude smaller 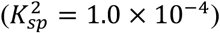 compared to the 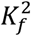 for a static phantom (e.g. 0.191), thus it can be neglected. Then s/p ratio is calculated from 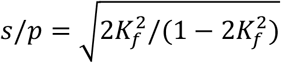 using the equation for unpolarized light^30^.

### Laser stability measurement

To test the stability during light steering, we used a photodiode (Thorlabs S121C) to measure the intensity at the distal end of each source fiber while maintaining constant voltage to the galvanometer controllers. The voltage applied to the two galvanometers controlled the mirrors’ positions, allowing manipulation of the beam direction along two dimensions (x and y). Light intensity was recorded using a power meter (Thorlabs PM400) and a Data Acquisition device (NI USB-6002). Light intensity time courses were obtained by averaging the photodiode output over a 4 ms window, repeated at 100 Hz. This matched the 4 ms exposure time and 100 Hz frame rate used for the SCOS cameras during human measurements. The measurement was repeated with a trapezoidal waveform galvanometer voltage input instead of a constant voltage, with upslope, plateau, downslope, and trough durations of 1.0, 5.2, 1.0, and 2.8 ms, respectively. The trapezoidal voltage waveform was also used to steer the beam between individual source fibers for human measurements. Intensity time courses were obtained by averaging over 4 ms windows, with each window starting 1 ms after the beginning of the plateau, as shown in Fig. 2a.

### Liquid dynamic phantom measurements

The dynamic phantom consists of an acrylic container filled with 1.6% Intralipid 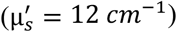^52^ at room temperature and a thin polytetrafluoroethylene tube (inner diameter = 0.082 inches, wall thickness = 0.005 inches) also containing the same Intralipid solution. The tube is held in place through two watertight holes in the acrylic container. The tube is connected to a syringe pump that enables the change of the flow rate within the tube. The fibers are arranged in a grid with the same SDS as used in human measurements (Fig. 1). This grid is rotated at three angles, 0^°^, 45^°^, and 90^°^ (Fig. 4), to demonstrate the system’s ability to observe a localized increase in *rD*_*B*_ − 1. At each angle, 400 seconds of measurement is taken. There are 6 blocks each consisting of 30s no-flow followed by 30s flow, ending with 40s no-flow to allow the *rD*_*B*_ − 1 time course to return to baseline as shown in Fig. 4a. A flow rate of 2 cm/s is used during the flow period to mimic biologically relevant flow changes. During no-flow, the syringe pump is turned off. To remove any temporal fluctuations (ΔOD or *rD*_*B*_ − 1) extraneous to the underlying medium, any principal component with similar coefficients (standard deviation/mean < 1) across channels sharing one source optode is regressed out. The six blocks are block-averaged to show the flow response.

### Participants

19 participants within aged 20 to 60 years, with no prior diagnosis or treatment of neurological disorders were recruited for this study. Sex, gender, race, and ethnicity were not considered during recruitment. Participants were recruited through on-campus advertisements including flyers. Of the 19 participants, four showed motion artifacts in intensity time courses in more than half the blocks, leaving 15 participants’ data for subsequent analysis. The experimental procedure and protocols were approved and carried out in accordance with the regulations of Institutional Review Board of Boston University. Each participant provided a signed written informed consent form prior to the experiment.

### Cap Placement

For SCOS human measurements, the same NinjaCap employed for fNIRS measurements is used^53^. The sources and detectors are placed in a NinjaCap, an in-house 3D printed fNIRS cap made of flexible material (NinjaFlex, NinjaTek) with NIRx grommets for compatibility with variable tension spring holders. The cap is placed on the head with respect to electroencephalogram (EEG)10–10 midline central electrode site (Cz), which is estimated through measurement of nasion, inion, left/right pre-auricular points^54,55^. The cap contains grommet holders arranged in the grid shown in Fig. 1d to ensure the proper placement of optodes. Optodes are placed inside of the grommets and held in place by a variable tension spring holder. A short baseline measurement is taken to confirm that there is a high enough photon flux for effective measurements. Hair parsing is done to move hair out from the light path to reduce light loss.

### Cardiac waveforms measurements

Cardiac pulsatile signal was recorded for all participants during the same experiment sessions in which they performed the WCS task (described below) by taking 20 seconds of baseline data before the task. This also served as an indication of successful cap and optode placement on the head.

### Word-Color Stroop task

Participants perform a WCS task containing two conditions (see Figure 5) based on the task used by Jahani *et al*^34^. The stimulus is presented via PsychoPy software^56^. For a given trial of the congruent condition, the participant is provided with two-word options and is instructed to select the word which has matching meaning and font color (e.g. the word “red” in red font color). For a given trial of the incongruent condition, the participant is provided three-word options in a row and a single word prompt below, none of which have self-matching meaning and font color. The participant is instructed to select the word option in the top row which has a meaning that matches the prompt word font color. Participants practice each condition before proceeding with the task. Trial durations are 3 s each, and a block has 6 consecutive trials of either all congruent or all incongruent conditions. A run has an initial 20 s rest period followed by 9 blocks of each condition for a total of 18 blocks, the order of which is randomized, with an inter-stimulus rest jittered between 25 to 30 seconds. The total time for one run is approximately 7 minutes.

### Human measurement post-processing

For human measurement, ΔOD and *rD*_*B*_ − 1 from short separation (8 mm) SDS channels is linearly regressed out from 19 mm and 33 mm SDS channel signals. The ΔOD and *rD*_*B*_ − 1 time series are filtered using a fifth-order Butterworth filter with a cutoff frequency of 0.2 Hz. The filtered data is block-averaged from -2 to 38 seconds with respect to the stimulus onset (defined as 0 second). This is done for both congruent and incongruent trials, resulting in a block-averaged activation time course for each trial type and each participant. Systemic physiology and superficial changes are removed using short separation regression. In short separation regression, the short separation channel data is linearly regressed out from the block-averaged time course. This is done for both ΔOD and *rD*_*B*_ − 1. The data is then averaged across all participants to acquire a group-averaged result. Since blocks contaminated by motion artifacts were excluded from analysis, we used a weighted average based on the number of valid blocks per subject for group averaging.

### Image reconstruction

To reconstruct brain activation from time series measured in all the source-detector channels, we follow the established methods for image reconstruction^43,49^, with the optical and dynamics properties of different brain layers listed in Supplemental Table 2. In short, we assume the data can be represented by a linear model that relates *rD*_*B*_ measured in the source-detector channels (denoted as *rD*_*B,SD*_) with *rD*_*B*_ in all the spatial locations on the head (denoted as 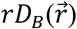)

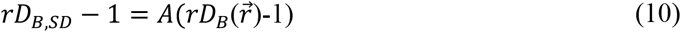

Where *A* is the sensitivity matrix. The sensitivity matrix was obtained using the head model simulations as previously described^49^, and the reconstruction is done by inverting Eq. 10 for the usual case of fewer number of measurements (*rD*_*B,SD*_) than unknowns 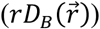,

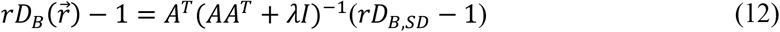

Here *A*^*T*^ is the transpose of *A*; *I* is the identity matrix; and *λ* = *αb* where *α* = 0.05 is the regularization parameter, and *b* is the largest eigenvalue of the matrix *AA*^*T*^. The image reconstruction for Δ*OD* is obtained similarly using already established functions in Cedalion^38^.

### Statistics

The Δ*OD* and *rD*_*B*_ − 1 responses are first averaged over all the blocks within each subject and then averaged over all the subjects to obtain the mean time course for each channel. The standard error across subjects are also obtained as shown by the shaded regions in Figs. 5c,d. Two-tailed t-tests were used to obtain the p value for the reported (mean±standard error) results, with the significance level set as 0.05. The mean, standard deviation and standard error were computed using built-in MATLAB functions.

## Acknowledgements

We thank Joseph O’Brien, De’Ja Rogers, Sudan Duwadi, Sreekanth Kura, Bingxue Liu, Allen Zhou, Lina Lin, Kyle Bohl, Matthew Simkulet, Zahid Yaqoob, and John T. Giblin for useful discussions. This project is supported by NIH UG3EB034710 (Cheng), NIH R01NS135081 (Franceschini).

## Author contribution

B.K., D.B., and X.C. conceived the project. B.K., A.H., E.H., B.Z., D.B., and X.C. prepared the electrical circuitry. B.K., A.H., B.Z., M.Robinson, M.Renna, S.C., M.F., D.B., and X.C. designed and built the optical system. B.K., A.H., and T.C. performed phantom preparation and measurement. B.K., A.H., J.A., and M.Y. designed and measured human cognitive task induced signals. B.K., A.H., and L.C. analyzed the data. B.K., A.H., D.B., and X.C. wrote the paper. All authors discussed the results and commented on the paper.

## Data Availability Statement

Data used to generate the main and supplementary figures are available on Zenodo (https://zenodo.org/records/15881287). All other datasets that support the findings of this study are available from the corresponding author upon reasonable request.

## Code availability statement

The SCOS control board code is written in C++ and the preprocessing and data analysis pipeline is written in MATLAB available on GitHub (https://github.com/BUNPC/2025-SCOS). The camera acquisition code is written in Python and available on GitHub (https://github.com/BUNPC/Multicamera-Imaging-Software). Image reconstruction is written in Python and available on GitHub (https://github.com/BUNPC/cedalion_scos-2025/blob/main/examples/head_models/imagerecon_scos.ipynb). All of the code has been deposited in Zenodo (https://zenodo.org/records/15857987). Any other code, including those used to generate the manuscript’s figures, are available from the corresponding author on reasonable request.

## Supplemental information

Supplemental Figures 1-2. Supplemental Tables 1-2.

