## Supplemental Figures and Tables for "Mapping human cerebral blood flow with high-density, multi-channel speckle contrast optical spectroscopy"

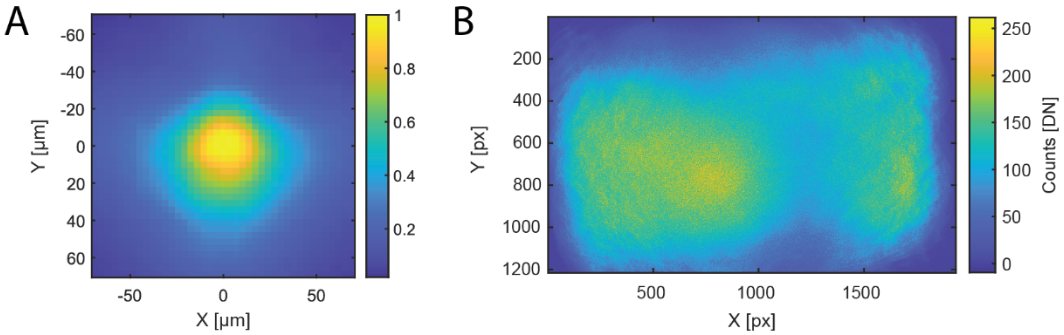


Supplemental Figure 1. (A) Example image of laser focal spot after focusing through the achromatic lens onto a camera. (B) An example of defocused image of the detector fiber bundle onto the camera, magnified with the 4f system.

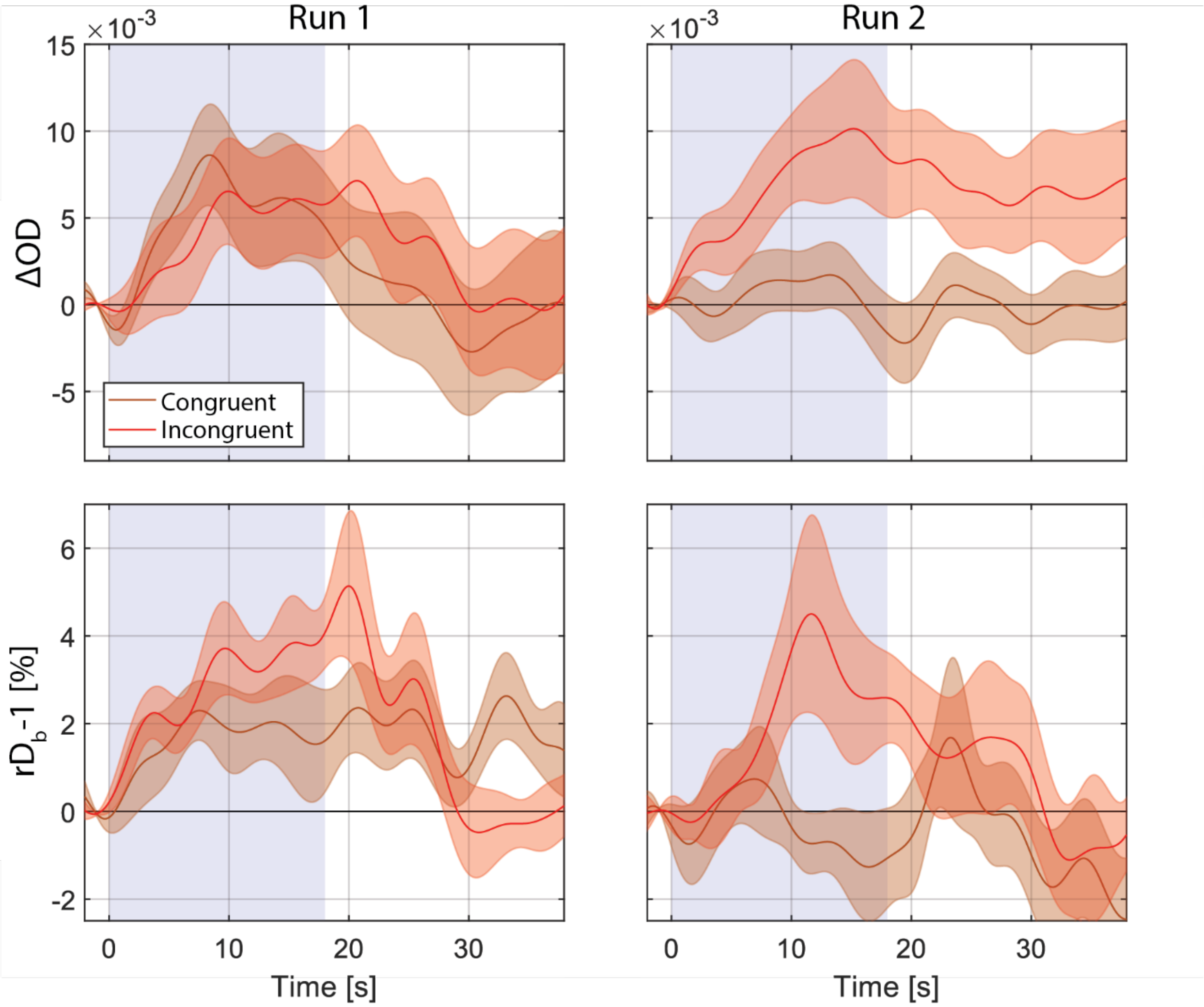


Supplemental Figure 2. Change in cognitive activation response over the two runs across participants, averaged across selected 33 mm SDS channels. The shaded area represents the standard error across subjects for the within-subject block-averaged time course.

| Subject | ΔOD | r$D_b - 1$ (%) |
|---|---|---|
| 1 | $2.7 \times 10^{-2}$ | 17 |
| 2 | $5.5 \times 10^{-2}$ | 21 |
| 3 | $6.8 \times 10^{-3}$ | 3.5 |
| 4 | $3.0 \times 10^{-2}$ | 9.0 |
| 5 | $2.1 \times 10^{-2}$ | 5.4 |
| 6 | $1.4 \times 10^{-2}$ | 4.4 |
| 7 | $1.0 \times 10^{-3}$ | 1.7 |
| 8 | $1.5 \times 10^{-2}$ | 5.7 |
| 9 | $4.1 \times 10^{-3}$ | 3.0 |
| 10 | $1.9 \times 10^{-2}$ | 5.3 |
| 11 | $1.8 \times 10^{-2}$ | 12 |
| 12 | $1.6 \times 10^{-2}$ | 5.3 |
| 13 | $1.1 \times 10^{-3}$ | 0.12 |
| 14 | $1.2 \times 10^{-2}$ | 3.8 |
| 15 | $1.8 \times 10^{-2}$ | 3.0 |

**Supplemental Table 1. Peak Channel Activation for each Subject.** Subject data was trial averaged, then temporally averaged from 10 – 15 seconds, the same range used for image reconstruction (Fig. 5). The peak activation in 33 mm SDS channels during the incongruent WCS task is shown for ΔOD and r$D_b - 1$. The average signal changes in the channel showing the largest activation were $1.7 \times 10^{-2}$ in ΔOD and 6.6% in r$D_b - 1$.

| | $\mu_a$ (1/mm) | $\mu_s'$ (1/mm) | $\alpha D_b$ ($mm^2/s$) |
|---|---|---|---|
| Skin | 0.0191 | 0.65934 | $1\times10^{-6}$ |
| Skull | 0.0136 | 0.85914 | $1\times10^{-6}$ |
| DM | 0.0191 | 0.65934 | $1\times10^{-6}$ |
| CSF | 0.0026 | 0.00999 | $1\times10^{-6}$ |
| GM | 0.0186 | 1.0989 | $6\times10^{-6}$ |
| WM | 0.0186 | 0.85914 | $6\times10^{-6}$ |

**Supplementary Table 2. Optical and Dynamic Properties of the Anatomical Head Model.** Table of the optical and dynamic properties for the anatomical head model used in Monte Carlo simulation representing the skin, skull, dura matter (DM), cerebrospinal fluid (CSF), gray matter (GM), and white matter (WM).
